# Targeting redox imbalance through Nrf2 activation in the inflamed coeliac duodenum

**DOI:** 10.64898/2026.04.22.720101

**Authors:** Heather Loughnane, Shrikanth Chomanahalli Basavarajappa, Anna Dominik, Conor Finlay, Seamus Hussey, Darren Ruane, Patrick T. Walsh

## Abstract

Coeliac Disease (CeD) is a chronic gastrointestinal inflammatory disease initiated by dietary gluten in genetically predisposed individuals. While the inflammatory processes which drive tissue destruction in the coeliac duodenum have been extensively characterised, an increased oxidative stress (OS) response has also been suggested to contribute to CeD pathogenesis. However, the precise mechanisms which regulate OS in the coeliac mucosa and whether they impact inflammation remain ill defined. The master anti-oxidant transcriptional regulator Nuclear factor erythroid 2-related factor 2 (Nrf2), and its inhibitor, Kelch like ECH-associated protein 1 (Keap1) have been implicated in chronic gastrointestinal inflammatory diseases, such as ulcerative colitis but have been largely unexplored in the context of CeD. To investigate redox balance in the CeD duodenum, we utilised single cell transcriptomics to assess overall OS and cytoprotective Nrf2 activation across cell subsets in duodenal biopsies from CeD patients. OS induced gene expression was broadly increased across multiple cell subsets in the CeD mucosa. Simultaneously, specific markers of Nrf2 activation were decreased in cell subtypes central to pathogenesis of CeD, including activated CD4^+^ T cells and intraepithelial T lymphocytes, indicating a distinct redox imbalance in these cells. Furthermore, pharmacological activation of Nrf2 significantly decreased gliadin induced *IFNG* expression in CeD duodenal biopsies. Taken together, our findings demonstrate that redox imbalance represents a therapeutic opportunity for the modulation of proinflammatory responses that drive the pathogenesis of CeD.

## Introduction

Coeliac disease (CeD) is an immune-mediated enteropathy triggered by dietary gluten in genetically susceptible individuals, leading to chronic inflammation and structural damage of the duodenal mucosa [1]. As a consequence of gluten induced immune activation, a complex network of cross-talk between stromal and immune cell subsets occurs which drives mucosal atrophy and villous blunting characteristic of CeD pathogenesis.

In the duodenum, partially digested gluten peptides are deamidated by tissue transglutaminase (tTG), enhancing their binding to HLA-DQ2/DQ8 molecules on antigen-presenting cells (APCs), and promoting the activation of gluten-specific CD4^+^ T cells in the lamina propria [2, 3]. This drives a T_H_1-skewed immune response characterised by the release of proinflammatory cytokines such as interferon-γ (IFN-γ) and interleukin (IL)-21, promoting downstream B cell activation and autoantibody production against tTG and deamidated gliadin peptides [3-5].

Concurrently, innate immune mechanisms amplify tissue damage. IL-15 produced by intestinal epithelial cells in response to stress, and proinflammatory cytokines released by gliadin-specific CD4^+^ T cells cooperate to give rise to highly proliferative intraepithelial T lymphocytes (T-IELs) with natural killer (NK)-like properties [6-8]. These cytotoxic T-IELs mediate epithelial cell apoptosis and contribute directly to villous atrophy, promoting tissue injury in the CeD duodenum [8, 9].

Oxidative Stress (OS) is characterised by an imbalance between the production of reactive oxygen and nitrogen species (ROS and RNS), and endogenous antioxidant defences, and is implicated in CeD pathogenesis. Several *in vitro* studies have demonstrated that gliadin exposure can induce OS in duodenal cells derived from CeD patients [10, 11]. Moreover, elevated plasma markers of OS in CeD patients have been reported to correlate with duodenal atrophy, indicating that OS functions in the pathogenesis of CeD [12, 13]. Furthermore, inflammatory cytokines and activated immune cells may exacerbate OS, creating a self-perpetuating cycle of tissue damage and impaired mucosal healing [14-16]. However, whether OS is dysregulated in the CeD duodenal mucosa, and if so, in which cell types, remains to be fully investigated.

To restrict OS induced tissue damage, endogenous antioxidant mechanisms act in a cytoprotective fashion. One such antioxidant defence pathway is mediated by Nuclear factor erythroid 2-related factor 2 (Nrf2), and its inhibitor, Kelch like ECH-associated protein 1 (Keap1) [17]. Mechanistically, Keap1 sequesters Nrf2 in the cytoplasm thereby inhibiting nuclear translocation and pathway activity. Increased ROS releases Keap1 from Nrf2, allowing Nrf2 nuclear entry and regulation of cytoprotective gene transcription through binding to Antioxidant Response Elements [17, 18]. While Nrf2 activity has emerged as an important regulator of mucosal barrier integrity and modulator of inflammation in the context of different gastrointestinal diseases such as ulcerative colitis, its function in the context of CeD is largely unexplored [19, 20]. It is noteworthy however that among its immunomodulatory functions, Nrf2 has been reported to act as a selective inhibitor of T_H_1 type responses, a key driver of CeD pathogenesis [21, 22].

To address some of these outstanding questions, we first investigated whether elevated OS is evident in the inflamed duodenum of CeD patients, utilising a single cell transcriptomic atlas we have recently generated [23]. Significantly heightened levels of OS were detected across multiple cell subsets of the inflamed duodenum in CeD. In contrast, the expression of a specific defined panel of Nrf2 dependent genes was found to be reduced in several cell types, including activated CD4^+^ T cells, and T-IEL subsets which contribute to elevated IFN-γ production and are central to mucosal tissue damage in CeD [23]. Based upon these findings, we assessed whether pharmacological activation of Nrf2 might alter pathogenic responses in the CeD mucosa. In this context, Nrf2 activation suppressed gliadin peptide induced *IFNG* expression in CeD duodenal explants while promoting antioxidant gene expression. Taken together, these data identify a specific redox imbalance in the CeD duodenal mucosa which may play an important orchestrating role in disease pathogenesis. Understanding the interplay between OS and immune responses in CeD may open new avenues for therapeutic interventions aimed at restoring redox homeostasis alongside strict adherence to a gluten-free diet.

## Methods

### Patient Information

Duodenal Biopsies from CeD and control patients were recruited as part of the Science Underlying CoeliaC Evolution: Explanations, Discoveries, and Solutions (SUCCEEDS) Study, at the gastroenterology unit at Our Ladys Childrens Hospital (Crumlin, Ireland), approved by the local Research Ethics Committee (REC-104-22). All biopsy samples were obtained at the point of diagnostic endoscopy, following informed consent/assent by individuals. All participants in the SUCCEEDS Study were consuming a full gluten-containing diet at the time of endoscopy. Cases were identified based on subsequent histological confirmation and Marsh scoring. Normal duodenal histology was confirmed in all control subjects. Patient demographics for the cohort used in *ex vivo* investigation can be found in **Table 1**. Information on patients included in the scRNAseq analysis can be found in our previous report [23].

**Table 1.**
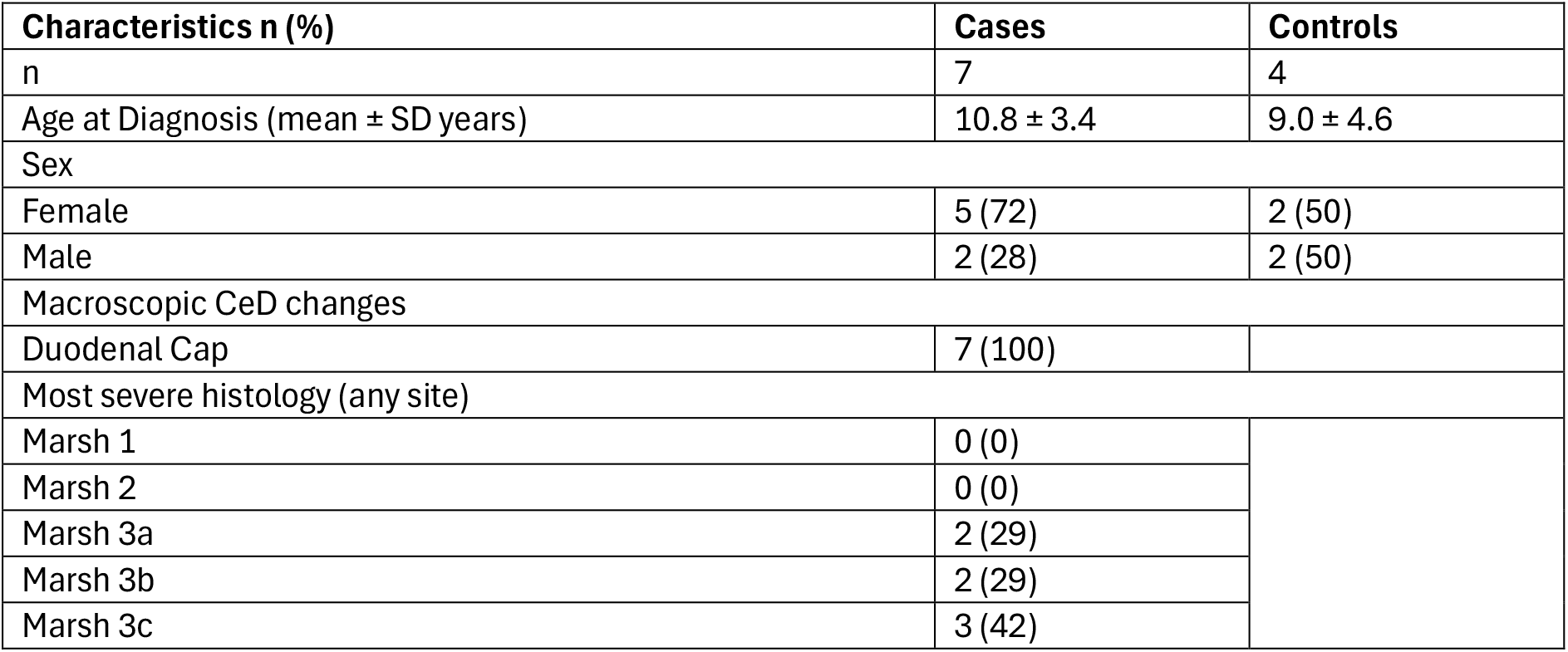
Patient Information for CeD duodenal biopsy explant studies. CeD, Coeliac Disease

### Data Acquisition

We utilised our previously processed single cell RNA sequence (scRNAseq) dataset, consisting of paediatric duodenal biopsies from 21 active CeD patients and 11 control patients, and can be accessed through the Gene Expression Omnibus (GEO) accession number, GSE277276 [23]. We used a recent bulk RNAseq dataset from adult duodenal biopsies obtained from 11 CeD patients and 5 control patients in our analysis and can be accessed through the GEO accession number, GSE146190 [24]. Gene sets to determine OS and Nrf2 responsive gene module scores were obtained from gene lists reported by Xu et al. and Morgenstern et al., respectively [25, 26].

### scRNAseq Data Analysis

The scRNAseq dataset was loaded into a Seurat Object in R for initial processing [27]. We utilised UCell to generate OS and Nrf2 activity module scores, and graphed module scores for each cell type and compartment using ggplot2 [28]. Information on initial data processing of the scRNAseq dataset used in analysis can be found in our previous report [23].

### Bulk RNAseq Data Analysis

Differential analysis was executed using DESeq2 [29]. Genes classified as differentially expressed matched the criteria of adjusted p-value < 0.01 and absolute log_2_Fold Change >0.5. Adjusted p-values were determined by the False Discovery Rate (FDR). Differential expression of genes was visualised using pheatmap.

### Duodenal Biopsy Stimulation

Duodenal biopsies isolated from paediatric CeD or control patients were either left unstimulated or stimulated with gliadin peptides (CSBio®, Shanghai, China) with or without Ki696 (MedChemExpress®, NJ, USA), in Dulbecco’s Modified Eagle’s Medium (Gibco™, MA, USA) supplemented with 10% fetal bovine serum (Corning, Inc., NY, USA), and 0.1% each of Penicillin-Streptomycin (Gibco™, MA, USA), Gentamycin B (Sigma-Aldrich, MO, USA), Amphotericin B (Gibco™, MA, USA), and HEPES (Sigma-Aldrich, MO, USA) for 18hrs at 37°C with 5% CO_2_. Information on gliadin peptides used in stimulation can be found in Table 2.

**Table 2.**
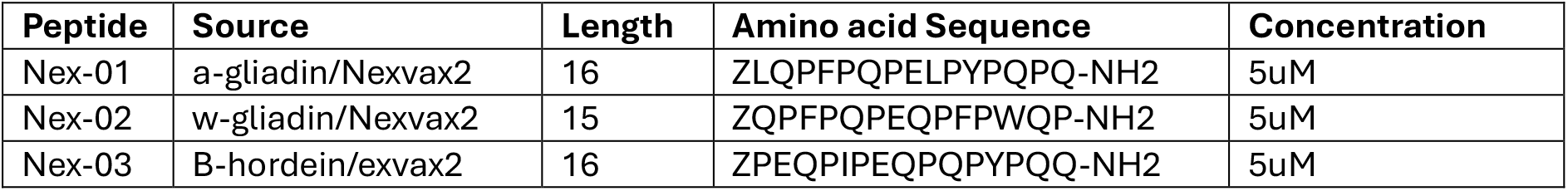
Gliadin Peptides used for Duodenal Biopsy Stimulation.

### RNA Isolation and Reverse Transcription

ISOLATE II RNA Mini Kit (Meridian Bioscience®, OH, USA) was used according to manufacturer’s instructions for RNA isolation from duodenal tissues. RNA was placed on ice once eluted and RNA concentration was measured using a DeNovix DS-11 nanodrop spectrometer (DeNovix®, DE, USA). All RNA samples were stored at −80°C for future use. RNA was reverse transcribed into cDNA using the High-Capacity cDNA Reverse Transcription Kit (Applied Biosystems™, MA, USA) according to manufacturer’s instructions and using a VeritiPro™ 96-well Thermal Cycler (Applied Biosystems™, MA, USA).

### Quantitative PCR

Quantitative polymerase chain reaction (qPCR) was performed with TaqMan™ Fast Advanced Master Mix (Applied Biosystems™, MA, USA) on a QuantStudio™ 3 (Applied Biosystems™, MA, USA). Cycle thresholds (CTs) were used to calculate the relative expression of target genes using 2^−Δ*CT*^. Primers used in this report are outlined in supplementary Table 3.

### Statistical Analysis

*Ex vivo* data analysis was performed using GraphPad Prism® version 10 (GraphPad Software, Inc., California, USA) and is presented as mean +/-s.e.m. According to normality, and whether data was paired, student t-test, and Mann-Whitney U test was applied for statistical analysis, as stated in figure legends

## Results

### Oxidative Stress is elevated in the CeD duodenal mucosa

To determine whether OS was elevated in the CeD duodenum, we investigated whether an OS gene expression signature was altered in specific cell subtypes using our previously described scRNAseq dataset [23]. OS module scores were generated for cell subsets using UCell based on a previously reported OS signature panel of 438 genes differentially expressed in the intestinal mucosa of Crohn’s Disease patients [26].

OS module scores were increased across broad cellular compartments in the CeD duodenum compared to control, with the exception of the epithelial cell compartment (Fig. 1a). At the cell subset level, OS gene expression signatures were elevated in multiple T cell subsets, including T-IEL subsets, CD4^+^ T cell subsets, T follicular helper cells and regulatory T cells (Tregs) (Fig. 1b) In addition, OS activation was also evident among myeloid cell subsets (DC1, monocyte-MNPs and macrophages) (Fig. 1c), B cells and plasma cells (Fig. 1d), enteroendocrine cells, and endothelial cells in the CeD duodenum compared to control (Fig. 1e,f). Collectively, these findings demonstrate that OS is broadly elevated across a multiple of cell types at sites of mucosal injury in the CeD duodenum.

**Figure 1.**
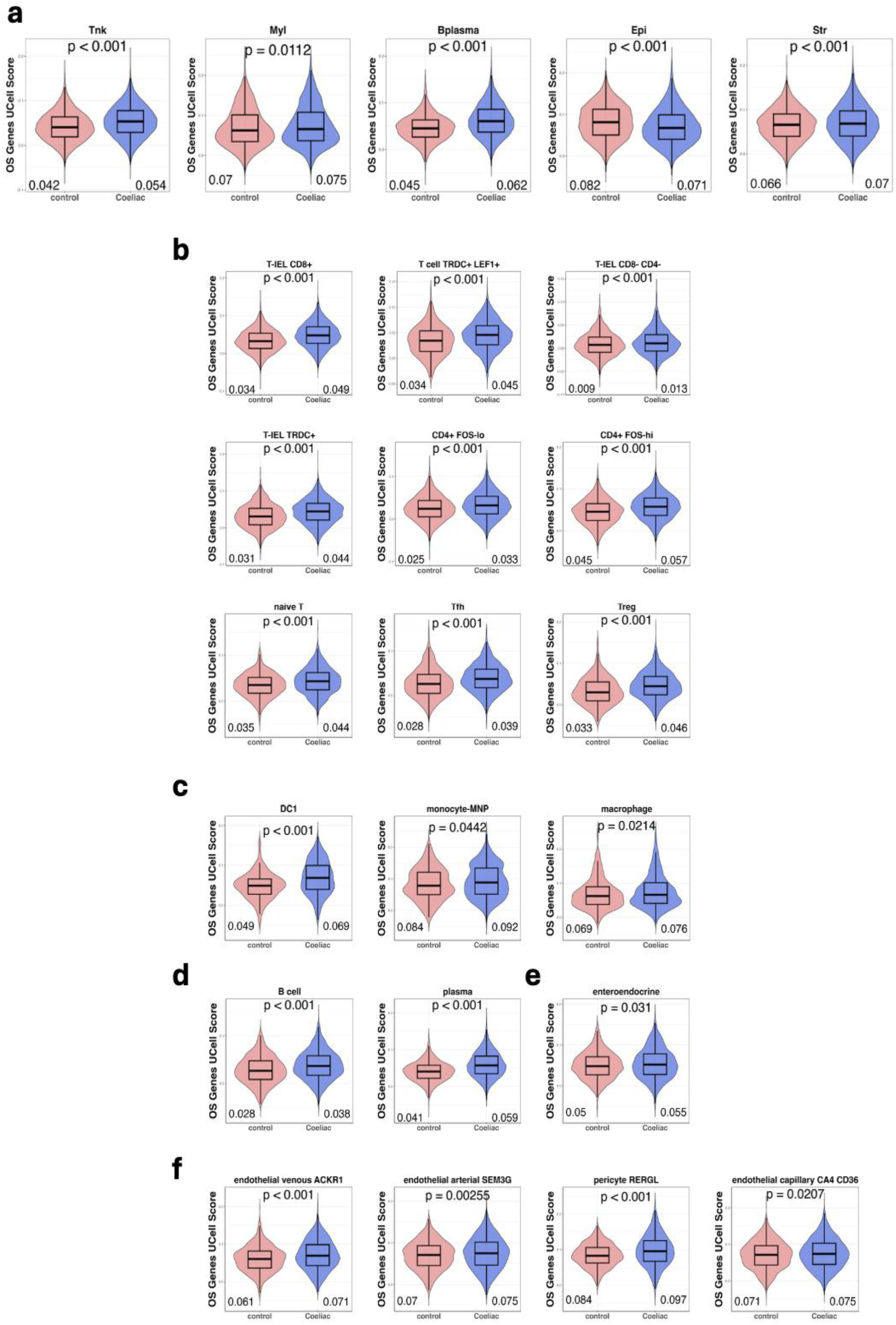
Oxidative Stress Module Scores are Increased across multiple cell types in CeD compared to control. Oxidative Stress (OS) related genes were stored as a module and OS module scores were generated using UCell. OS module scores between CeD and control were compared by cellular compartment (a), and OS module scores for each cell type were generated for cells within the T and NK (b), myeloid (c), B plasma (d), epithelial (e) and stromal compartments (f). Cell types shown have significantly increased OS-module scores in CeD compared to control. OS, Oxidative Stress. Statistical analysis performed using Mann-Whitney U test.

### Nrf2 induced gene expression is decreased in the CeD duodenum

A redox imbalance between damaging prooxidant and antioxidant responses is an established characteristic of chronic mucosal inflammatory conditions [12, 13, 30, 31]. Given the broad increase in OS across multiple cell types in the CeD duodenum, we next investigated whether the activity of the antioxidant master transcriptional regulator, Nrf2 is similarly altered in the CeD mucosa. Nrf2 is constitutively expressed but post-transcriptionally regulated by Keap1 [17]. Thus, to assess Nrf2 activity, we employed a validated panel of established downstream Nrf2-dependent genes, determined by Morganstern et al. (2024) [25].

We first analysed a publicly available bulk RNAseq dataset (GSE146190), consisting of duodenal biopsies from adult CeD patients and control patients. We analysed relative expression levels of genes previously implicated in mucosal inflammation in CeD as well as expression levels of, *NFE2L2* (Nrf2), *KEAP1*, and the robust panel of Nrf2 activation marker genes, described above [25]. As expected, CeD duodenal biopsies exhibited increased *IL15, IFNG* and *KLRK1* (NKG2D) expression compared to control biopsies (Fig. 2a). Interestingly, and in contrast with the elevated OS signature, expression levels of *NFE2L2, KEAP1*, and established Nrf2 activation markers (*NQO1, GCLC, GCLM, HMOX1* and *TXNRD1*) were significantly downregulated in the CeD mucosa compared to controls (Fig. 2a).

**Figure 2.**
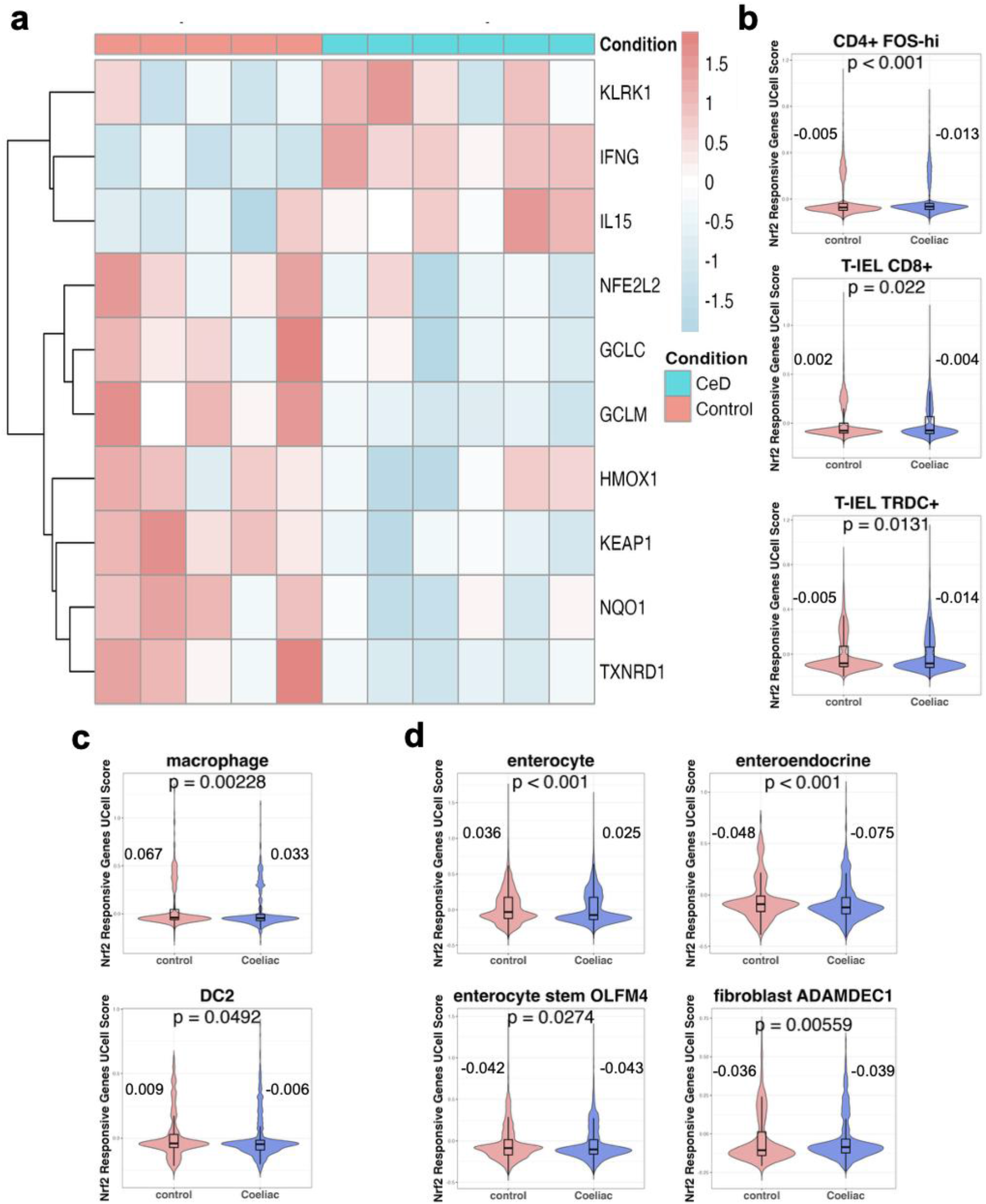
Nrf2 is downregulated at the global transcriptional level, and in cell types central to CeD pathogenesis. Expression of CeD related genes and Nrf2 activity related genes was determined for GSE146190, consisting of bulk RNAseq data from duodenal biopsies from HC (n=5) and CeD (n=11) (a). Nrf2 activity responsive genes were stored as a module and UCell module scores for Nrf2 responsive genes were generated for cells in our scRNAseq dataset. Nrf2 responsive gene module scores of CeD and controls were compared for cells within the T and NK (b), myeloid (c) and epithelial (d) compartment. Cell types with significantly reduced Nrf2 module scores in CeD compared to control are shown. Mean UCell values for control and CeD are annotated. Statistical analysis was performed by Mann-Whitney U test.

Given this imbalance between prooxidant and antioxidant responses, we next investigated whether reduced Nrf2 activity was evident in specific cell subsets in the CeD duodenum. To address this, UCell module scores using the Nrf2 activity gene panel were generated across all identified cell types in our scRNAseq dataset [25]. Intriguingly, while most cell types from CeD patients displayed no decrease in Nrf2 activity (Supp. Fig. 1), activated CD4^+^ T cells (CD4 FOS^hi^), T-IEL CD8^+^ and T-IEL TRDC^+^ cells from the duodenum of CeD patients had reduced Nrf2 activity module scores when compared to control patients (Fig. 2b). Similarly, macrophages, DC2 cells, enterocytes, enteroendocrine, tuft, enterocyte stem OLFM4^+^ and fibroblast ADAMDEC^+^ cells also had reduced Nrf2 activity signature scores in CeD patients compared to controls (Fig. 2c, d).

These data demonstrate a redox imbalance in the CeD duodenum, characterised by elevated OS, and reduced Nrf2 activation across several cell subsets central to CeD pathogenesis.

### Pharmacological activation of Nrf2 inhibits gliadin peptide induced inflammation in the CeD duodenum

Having established that redox imbalance is characteristic of the CeD duodenal mucosa and that Nrf2 activity is reduced in several cell types that play central roles CeD pathogenesis, we next investigated whether enhancing Nrf2 activation in the CeD mucosa may offer a therapeutic opportunity. To address this, we analysed the effects of a Keap1 inhibitor tool compound, Ki696, in CeD duodenal biopsies stimulated with gliadin peptides to elicit a tissue antigen specific response which models the effects of gluten exposure *in vivo*.

Gliadin peptide stimulation significantly increased *IFNG* expression, while significantly decreasing the expression of Nrf2 activation marker, *GCLC* in CeD duodenal biopsies (Fig. 3a, b). Treatment with Ki696 restored Nrf2 activity in CeD biopsies, evidenced through the induction of the Nrf2 target genes, *NQO1* and *GCLC* (Fig. 3b, c). Strikingly, Ki696 significantly attenuated *IFNG* expression in CeD biopsies stimulated with gliadin peptides, indicating that restoring Nrf2 activity and redox imbalance can modulate a key pathogenic pathway in CeD (Fig. 3a). These findings provide initial translational validation to support the targeting of redox pathways as a potential treatment for CeD patients.

**Figure 3.**
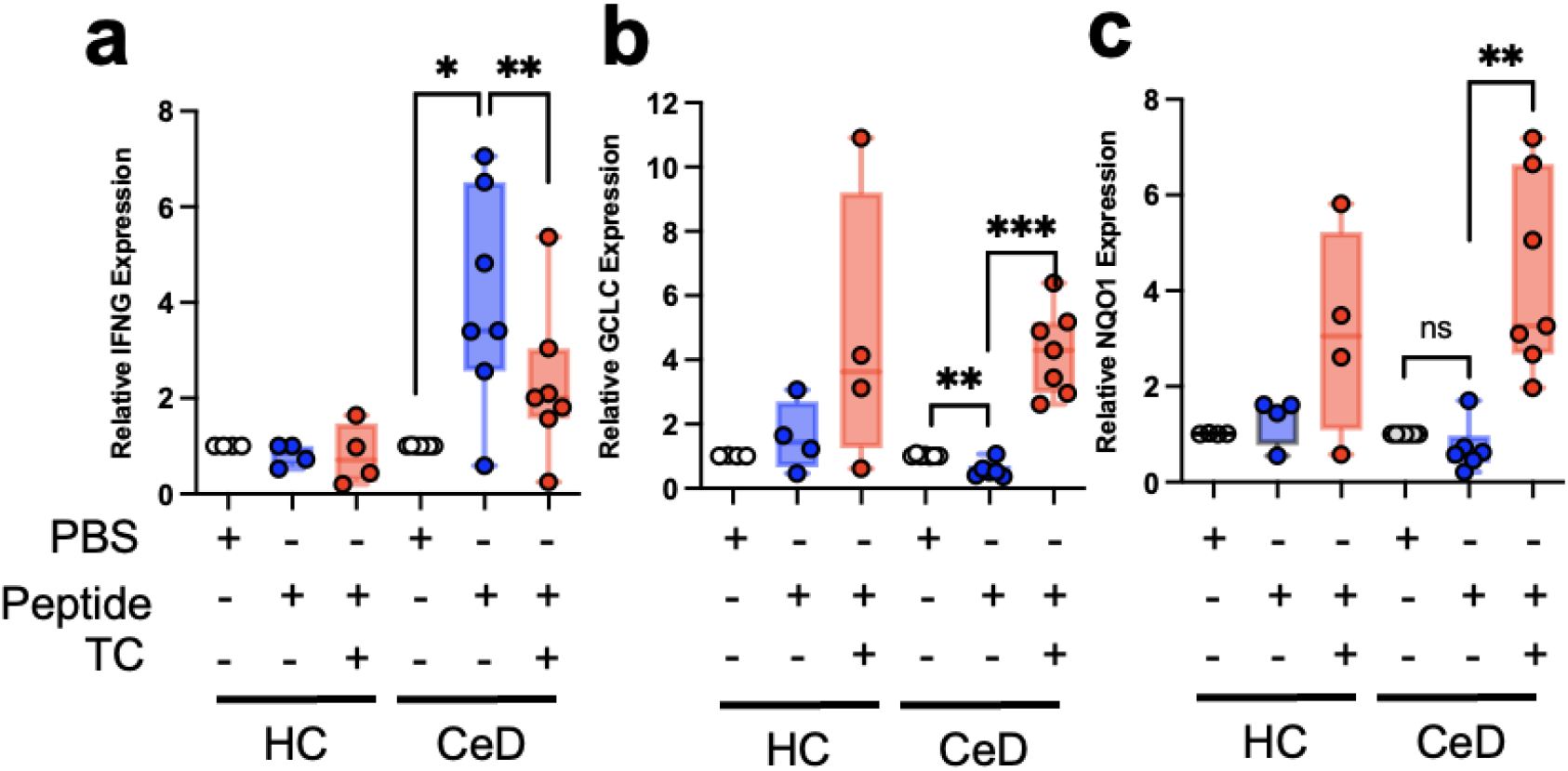
Activation of Nrf2 inhibits gluten mediated expression of *IFNG* in CeD biopsies *ex vivo*. Duodenal biopsies from individuals with CeD (n=7) or healthy control (n=4) were stimulated with or without gliadin peptides in the presence or absence of Ki696 (TC) for 18hrs, and subsequently analysed for *IFNG* (a), *GCLC* (b) and *NQO1* (c) mRNA expression by qPCR. Statistical analysis was performed by paired student t-test.

## Discussion

CeD is a chronic immune mediated enteropathy of the small intestine triggered by exposure to dietary gluten. The pathogenic mechanisms which drive antigen specific mucosal tissue destruction in the duodenum of CeD patients involves antigen deamidation and subsequent autoactivation of CD4+ T helper cells which orchestrate mucosal inflammation. Although, OS activation has been implicated in these events, the status of the OS response across distinct cell subsets has not been established [10-13]. Moreover, whether OS can impact pathogenic inflammation has not been addressed.

We have demonstrated a global increase in the OS transcriptional response in the duodenum of CeD patients compared to control patients. These data are consistent with previous reports of elevated OS markers in CeD duodenal biopsies and serum [12, 30-32]. Using scRNAseq analysis, we have further demonstrated that the OS signature is increased across the majority of major cell types in the CeD mucosa, indicating that elevated OS is evident across several cell compartments (Fig. 1). These findings prompted us to investigate whether regulation of OS may be impaired in this setting.

As Nrf2 is established as a master regulator of the antioxidant response, and is known to regulate mucosal inflammation in the large intestine [19,20], we asked whether Nrf2 activity might be similarly altered across cell subsets. Firstly, we demonstrated, using bulk transcriptomic analysis, that in contrast to broad OS activation, a robust panel of Nrf2 gene expression biomarkers were decreased in the CeD duodenum. A deeper analysis at the single cell level demonstrated that this decrease was evident across distinct cell subsets, including proinflammatory T cells as well as myeloid and epithelial cell types (Fig.2) Importantly, decreased Nrf2 activity occurred despite elevated OS activation among these cells. These data indicate a specific redox imbalance, between pathogenic OS activity and cytoprotective Nrf2 dependent anti-oxidant responses, among distinct cellular compartments which may contribute to disease pathogenesis.

Significantly, this redox imbalance was evident in activated CD4^+^ FOShi T cells, as well as γδT-IELs and CD8^+^ T -IELs subsets in the inflamed mucosa. Activated gliadin-specific CD4^+^ T cells are well established as central to CeD pathogenesis through the production of proinflammatory cytokines, such as IFN-γ [4, 37]. Further, gliadin-specific CD4^+^ T cells cooperate with IL-15, produced by stress-induced intestinal epithelial cells to promote NK-like receptor expression, such as NKG2D, in T-IELs to generate proinflammatory T-IELs which mediate intestinal epithelial injury in CeD [6-8, 23, 38]. Notably, we, and others, have previously identified γδT-IELs and CD8^+^ T-IELs, as well as CD4+ T cells, as major sources of *IFNG* expression in the CeD duodenum [6, 23]. Together, these findings suggest that impaired Nrf2 activity occurs in IFN-γ producing cells that are central to CeD pathogenesis, and may contribute to proinflammatory signalling and mucosal injury.

To further assess the *ex vivo* effects of Nrf2 activation on gluten induced inflammation, we stimulated CeD duodenal biopsy explants with gliadin peptides in the presence or absence of the Keap1 inhibitor, Ki696. As expected, the addition of Ki696 led to a robust induction of the Nrf2 responsive genes *GCLC* and *NQO1* in this setting. Interestingly, the gliadin peptides used in this study function specifically to activate HLA-DQ2.5^+^ gliadin-specific CD4^+^ T cells and the reduction in *GCLC* expression observed in CeD biopsies stimulated in the absence of Ki696, suggests that antigen-specific CD4^+^ T cell activation alone impaired the Nrf2 transcriptional activity in duodenal CeD biopsies.

Strikingly, inhibition of Keap1 reduced gliadin-induced *IFNG* expression in CeD duodenal biopsies. Having demonstrated that Nrf2 transcriptional activity is reduced in principal IFN-γ producing populations, specifically activated CD4^+^ T cells, CD8^+^ T-IELs and γδT-IELs in the CeD duodenum, these findings suggest that restoration of Nrf2 activity dampened *IFNG* expression in these cells in response to gliadin stimulation. It is unclear whether these effects occur specifically in the T cell subsets described, or as a result of the induction of Nrf2 activation among other cell types in the mucosal tissue. Indeed, our analysis has also demonstrated that Nrf2 activity is reduced among other cell types, most notably dendritic cells and myeloid cell subsets. However, it is also noteworthy that Nrf2 activation has previously been reported to directly suppress IFNγ expression under Th1 differentiation conditions *in vitro* [21].

Collectively, these data demonstrate the breadth of OS activation across distinct cell subsets in the inflamed CeD duodenum and provide evidence for a distinct redox imbalance across key pathogenic cell subsets. Importantly, this data also indicates that a restoration of Nrf2 activation can reduce proinflammatory processes central to CeD pathogenesis.

## Figure Legends

**Supplementary Figure 1.**
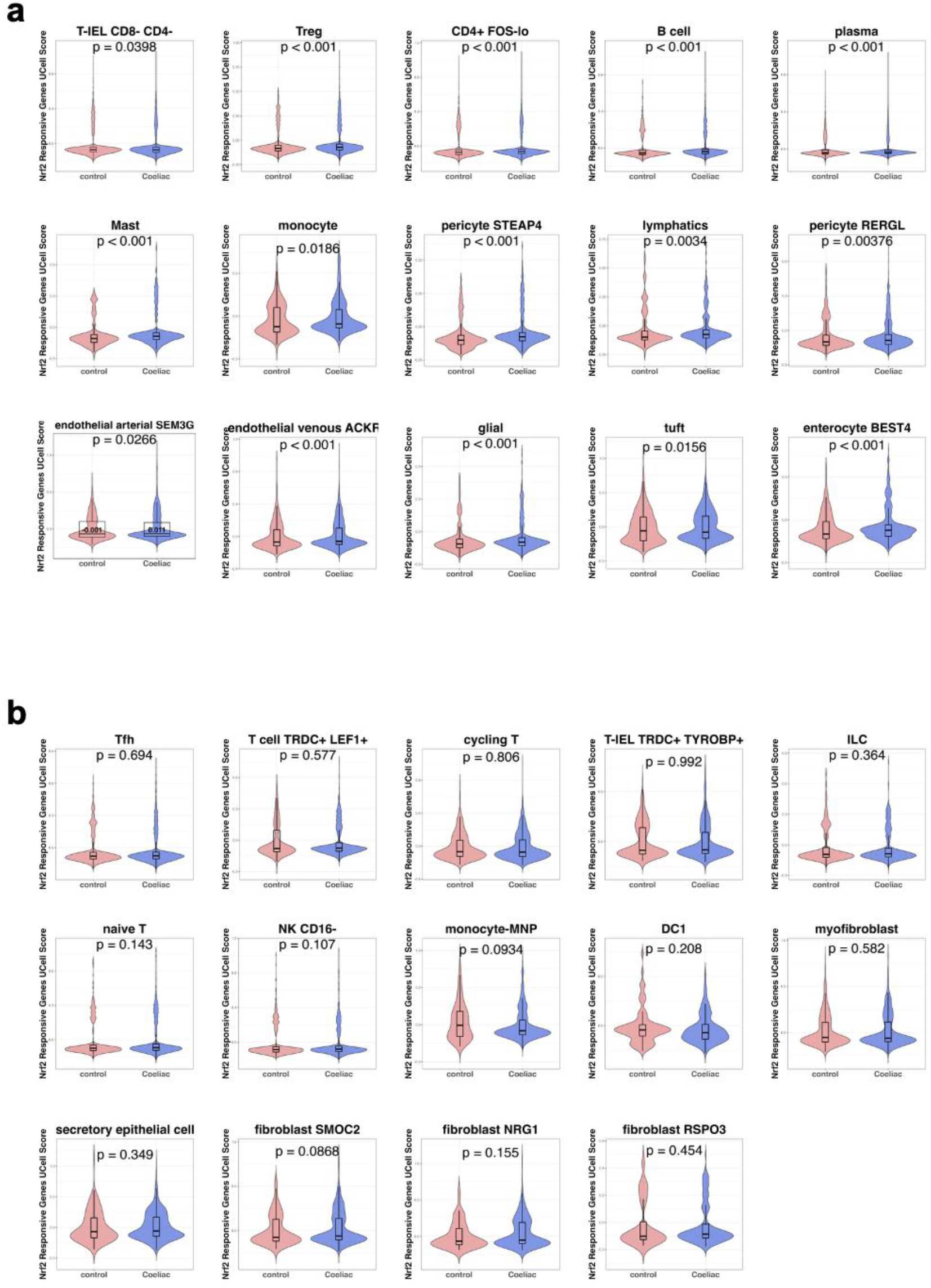
Nrf2 Activation Module Scores in Coeliac Disease vs Control. Nrf2 activity responsive genes were stored as a module and UCell module scores for Nrf2 responsive genes were generated for cells in our scRNAseq dataset. Nrf2 responsive gene module scores of CeD and controls were compared, for cell types where (a) Coeliac Disease cells had increased module scores, and (b)differences in module scores did not reach significance. *Statistical analysis was performed by Mann-Whitney U test*.

**STable 1.** Primers used for Quantitative PCR.

| Gene | Assay ID |
| --- | --- |
| NQO1 | Hs01045993_g1 |
| GCLC | Hs00155249_m1 |
| *IFNG* | Hs00989291_m1 |

## References

1. Green, P.H., B. Lebwohl, and R. Greywoode, Celiac disease. J Allergy Clin Immunol, 2015. 135(5): p. 1099–106; quiz 1107.

2. Hausch, F., et al., Intestinal digestive resistance of immunodominant gliadin peptides. Am J Physiol Gastrointest Liver Physiol, 2002. 283(4): p. G996–G1003.

3. Molberg, O., et al., Tissue transglutaminase selectively modifies gliadin peptides that are recognized by gut-derived T cells in celiac disease. Nat Med, 1998. 4(6): p. 713–7.

4. Nilsen, E.M., et al., Gluten specific, HLA-DQ restricted T cells from coeliac mucosa produce cytokines with Th1 or Th0 profile dominated by interferon gamma. Gut, 1995. 37(6): p. 766–76.

5. Bodd, M., et al., HLA-DQ2-restricted gluten-reactive T cells produce IL-21 but not IL-17 or IL-22. Mucosal Immunol, 2010. 3(6): p. 594–601.

6. Di Sabatino, A., et al., Epithelium derived interleukin 15 regulates intraepithelial lymphocyte Th1 cytokine production, cytotoxicity, and survival in coeliac disease. Gut, 2006. 55(4): p. 469–77.

7. Meresse, B., et al., Coordinated induction by IL15 of a TCR-independent NKG2D signaling pathway converts CTL into lymphokine-activated killer cells in celiac disease. Immunity, 2004. 21(3): p. 357–66.

8. Korneychuk, N., et al., Interleukin 15 and CD4(+) T cells cooperate to promote small intestinal enteropathy in response to dietary antigen. Gastroenterology, 2014. 146(4): p. 1017–27.

9. Hue, S., et al., A direct role for NKG2D/MICA interaction in villous atrophy during celiac disease. Immunity, 2004. 21(3): p. 367–77.

10. Luciani, A., et al., Lysosomal accumulation of gliadin p31-43 peptide induces oxidative stress and tissue transglutaminase-mediated PPARgamma downregulation in intestinal epithelial cells and coeliac mucosa. Gut, 2010. 59(3): p. 311–9.

11. Dolfini, E., et al., Damaging effects of gliadin on three-dimensional cell culture model. World J Gastroenterol, 2005. 11(38): p. 5973–7.

12. Boda, M., I. Nemeth, and D. Boda, The caffeine metabolic ratio as an index of xanthine oxidase activity in clinically active and silent celiac patients. J Pediatr Gastroenterol Nutr, 1999. 29(5): p. 546–50.

13. Stojiljkovic, V., et al., Glutathione redox cycle in small intestinal mucosa and peripheral blood of pediatric celiac disease patients. An Acad Bras Cienc, 2012. 84(1): p. 175–84.

14. Maluf, S.W., et al., DNA damage, oxidative stress, and inflammation in children with celiac disease. Genet Mol Biol, 2020. 43(2): p. e20180390.

15. Rivabene, R., E. Mancini, and M. De Vincenzi, In vitro cytotoxic effect of wheat gliadin-derived peptides on the Caco-2 intestinal cell line is associated with intracellular oxidative imbalance: implications for coeliac disease. Biochim Biophys Acta, 1999. 1453(1): p. 152–60.

16. Yu, T., et al., The Nutritional Intervention of Resveratrol Can Effectively Alleviate the Intestinal Inflammation Associated With Celiac Disease Induced by Wheat Gluten. Front Immunol, 2022. 13: p. 878186.

17. Bellezza, I., et al., Nrf2-Keap1 signaling in oxidative and reductive stress. Biochim Biophys Acta Mol Cell Res, 2018. 1865(5): p. 721–733.

18. Itoh, K., et al., Regulatory mechanisms of cellular response to oxidative stress. Free Radic Res, 1999. 31(4): p. 319–24.

19. Chen, H., et al., Butyrate ameliorated ferroptosis in ulcerative colitis through modulating Nrf2/GPX4 signal pathway and improving intestinal barrier. Biochim Biophys Acta Mol Basis Dis, 2024. 1870(2): p. 166984.

20. Peng, S., et al., The role of Nrf2 in the pathogenesis and treatment of ulcerative colitis. Front Immunol, 2023. 14: p. 1200111.

21. Turley, A.E., J.W. Zagorski, and C.E. Rockwell, The Nrf2 activator tBHQ inhibits T cell activation of primary human CD4 T cells. Cytokine, 2015. 71(2): p. 289–95.

22. Rockwell, C.E., et al., Th2 skewing by activation of Nrf2 in CD4(+) T cells. J Immunol, 2012. 188(4): p. 1630–7.

23. Richards, D., et al., Immune signaling mediates stromal changes to support epithelial reprogramming in celiac duodenum. Cell Rep, 2025. 44(8): p. 116039.

24. van der Graaf, A., et al., Systematic Prioritization of Candidate Genes in Disease Loci Identifies TRAFD1 as a Master Regulator of IFNgamma Signaling in Celiac Disease. Front Genet, 2020. 11: p. 562434.

25. Morgenstern, C., et al., Biomarkers of NRF2 signalling: Current status and future challenges. Redox Biol, 2024. 72: p. 103134.

26. Xu, S., et al., Oxidative stress gene expression, DNA methylation, and gut microbiota interaction trigger Crohn’s disease: a multi-omics Mendelian randomization study. BMC Med, 2023. 21(1): p. 179.

27. Hao, Y., et al., Dictionary learning for integrative, multimodal and scalable singlecell analysis. Nat Biotechnol, 2024. 42(2): p. 293–304.

28. Andreatta, M. and S.J. Carmona, UCell: Robust and scalable single-cell gene signature scoring. Comput Struct Biotechnol J, 2021. 19: p. 3796–3798.

29. Love, M.I., W. Huber, and S. Anders, Moderated estimation of fold change and dispersion for RNA-seq data with DESeq2. Genome Biol, 2014. 15(12): p. 550.

30. Daniels, I., et al., Elevated expression of iNOS mRNA and protein in coeliac disease. Clin Chim Acta, 2005. 356(1-2): p. 134–42.

31. Stojiljkovic, V., et al., Antioxidant status and lipid peroxidation in small intestinal mucosa of children with celiac disease. Clin Biochem, 2009. 42(13-14): p. 1431-7.

32. Ferretti, G., et al., Celiac disease, inflammation and oxidative damage: a nutrigenetic approach. Nutrients, 2012. 4(4): p. 243–57.

33. Cheng, J., et al., Duodenal microbiota composition and mucosal homeostasis in pediatric celiac disease. BMC Gastroenterol, 2013. 13: p. 113.

34. Mention, J.J., et al., Interleukin 15: a key to disrupted intraepithelial lymphocyte homeostasis and lymphomagenesis in celiac disease. Gastroenterology, 2003. 125(3): p. 730–45.

35. Iversen, R. and L.M. Sollid, The Immunobiology and Pathogenesis of Celiac Disease. Annu Rev Pathol, 2023. 18: p. 47–70.

36. Torinsson Naluai, A., et al., Whole genome transcriptional analysis of intestinal biopsies and blood cells indicate genes involved in antioxidant defense systems, amino acid metabolism and antigen presentation in the pathogenesis of celiac disease. BMC Med, 2025. 23(1): p. 507.

37. Risnes, L.F., et al., Disease-driving CD4+ T cell clonotypes persist for decades in celiac disease. J Clin Invest, 2018. 128(6): p. 2642–2650.

38. Tang, F., et al., Cytosolic PLA2 is required for CTL-mediated immunopathology of celiac disease via NKG2D and IL-15. J Exp Med, 2009. 206(3): p. 707–19.

